# Circadian Oscillations Persist in Low Malignancy Breast Cancer Cells

**DOI:** 10.1101/664664

**Authors:** Sujeewa S. Lellupitiyage Don, Hui-Hsien Lin, Jessica J. Furtado, Maan Qraitem, Stephanie R. Taylor, Michelle E. Farkas

## Abstract

Epidemiological studies have shown that humans with altered circadian rhythms have higher cancer incidence, with breast cancer being one of the most cited examples. To uncover how circadian disruptions may be correlated with breast cancer and its development, prior studies have assessed the expression of *BMAL1* and *PER2* core clock genes via RT-qPCR and western blot analyses. These and our own low-resolution data show that *BMAL1* and *PER2* expression are suppressed and arrhythmic. We hypothesized that oscillations persist in breast cancer cells, but due to limitations of protocols utilized, cannot be observed. This is especially true where dynamic changes may be subtle. In the present work, we generated luciferase reporter cell lines representing high- and low-grade breast cancers to assess circadian rhythms. We tracked signals for *BMAL1* and *PER2* to determine whether and to what extent oscillations exist and provide initial correlations of circadian rhythm alterations with breast cancer aggression. In contrast to previous studies, where the clock was deemed to be “broken” in breast cancer, our luminometry data reveal that circadian oscillations of *BMAL1* and *PER2* do in fact exist in the low-grade, luminal A cell line, MCF7 but are not apparent in high-grade, basal MDA-MB-231 cells. To our knowledge, this is the first evidence of core circadian clock oscillations in breast cancer cells. This work also suggests that circadian rhythms are increasingly disrupted with breast cancer tumor grade/aggressiveness, and that use of real time luminometry to study additional representatives of breast and other cancer subtypes is highly merited.

## Introduction

Circadian rhythms are biological changes that follow a daily cycle, responding primarily to light and darkness in an organism’s environment. Their disruption has been associated with increased risk of several cancers, including breast.^1^ Circadian rhythms are regulated by a molecular circadian clock comprising CLOCK, BMAL, PER, and CRY proteins. Oscillations of these and ancillary circadian genes have been found to be arrhythmic/absent in several disease models and tissues, including peripheral blood mononuclear cells from chronic myeloid leukemia,^2^ colorectal liver metastases,^3^ and prostate^4^ and breast cancer cell lines.^5–8^ In most of these cases, RT-qPCR, western blots, and immunohistochemistry were used to evaluate circadian changes. These assays result in less-detailed oscillation assessments because sampling frequency is typically 4-6 h and the experiment covers ∼48 h; more time-points and collection over a longer duration are difficult to achieve. Similar studies revealed that core clock gene transcripts are present, but do not oscillate in breast cancer cell lines.^6–10^ As examples, in work using MCF7 and MDA-MB-231 (among other breast cancer) cells, RT-qPCR and DNA/miRNA microarray data show that core circadian genes *BMAL1* and *PER2* lack rhythmic oscillations, although some clock controlled genes (CCGs) and miRNAs are rhythmic.^6,8^ Another study in MCF7 cells indicated that upon synchronization, some core clock genes may show fluctuations, but not all (including *PER1* and *PER2*).^7^ Taken together, since these findings were based on RT-PCR, DNA/miRNA micro-array, and Western blotting data, we considered that further, more detailed investigation was merited. Using these methods it is difficult to separate trends from background, especially in cases of subtle changes and low amplitudes, resulting in reduced accuracy of period and rhythmicity evaluations. To determine whether clocks oscillate in cancer, continuous tracking in living cells is needed. The use of reporter systems combined with luminometry can facilitate tracking of circadian oscillations at a higher resolution via data recording at shorter intervals over a longer period.^11^ Increased data collection enables more accurate analyses of circadian parameters. While luminometry has been used to track circadian rhythms in plants, cyanobacteria, mice, and human cells,^11^ this work is the first instance of its use for tracking circadian rhythms in breast cancer models.

According to the 2017 American Cancer Society annual report, breast cancer is the most common cancer (30% of estimated new cancer cases) among women in the United States.^12^ Breast cancers may be divided into four molecular subtypes based on gene expression profile: luminal A, luminal B, HER2+, and basal (or triple-negative).^13^ Tumor grade and aggressiveness increase from luminal A to basal subtypes. We hypothesized that RT-qPCR-based analyses are insufficient to track circadian oscillations of breast cancer cells. Furthermore, as circadian disruptions have been associated with disease severity, we hypothesized that with increasing breast tumor grade/aggressiveness, cellular circadian rhythms are increasingly disrupted. To test these hypotheses, we used RT-qPCR and luminometry to track *BMAL1* and *PER2* transcription and western blots to confirm their levels of expression in two breast cancer cell lines, MCF7 (luminal A) and MDA-MB-231 (basal). By using the higher resolution luminometry technique, we find that circadian oscillations exist in low grade, luminal A MCF7 breast cancer cells, but not high grade, basal MDA-MB-231 cells.

## Materials and Methods

### Cell Culture

HEK293T and MCF7 cells were obtained from Prof. D. Joseph Jerry (Veterinary and Animal Sciences, UMass Amherst). MCF7s were characterized for expression of estrogen receptor and its signaling; both were mycoplasma tested prior to transfer via PCR Mycoplasma detection kit (AMB). MDA-MB-231 cells were obtained from Prof. Shelly Peyton (Chemical Engineering, UMass Amherst), which were evaluated for karyotype and mycoplasma, along with subtype confirmation via RNAseq prior to transfer. HEK293T cells were maintained in DMEM:F12 supplemented with 10% FBS, 1% penicillin-streptomycin, and 0.015 mg/mL gentamicin. MCF7 and MDA-MB-231 cells were maintained in DMEM, supplemented with 5% FBS, 1% penicillin-streptomycin, and 1% L-glutamine. All media and supplements were from Gibco. Cells were incubated at 37°C under 5% CO_2_ atmosphere.

### Synchronization of Cells by Serum Shock

Cells were seeded in 35 mm culture dishes at 2 × 10^5^ cells/mL and incubated for 1-3 days. Confluent cells were washed with PBS (Gibco) and starved in DMEM media supplemented only with 1% L-glutamine for 12 h. Cells were then serum shocked using DMEM containing 50% FBS and 1% L-glutamine for 2 h.

### RNA Extraction and cDNA Synthesis

Following synchronization, cells were washed with PBS and returned to starvation conditions. Cells were harvested with the first time point (T=0) taken prior to serum shock, and every 4 h thereafter for 48 h. Total RNA was extracted via TRIzol Reagent (Gibco) according to manufacturer’s instructions (detailed protocols in Supporting Information).

### Quantitative Real-Time PCR (RT-qPCR)

RT-qPCR was performed in 96-well plates. The reaction consisted of 100 ng cDNA, 10 µL iTaq universal SYBR Green Supermix (Biorad), 4 µM each forward and reverse primer (Integrated DNA Technologies), and RNAse-free water to 20 µL. Primer sequences are in Supporting Information. After brief centrifugation, samples were analyzed via CFX Connect real-time system (Biorad) programmed with an initial denaturation at 95°C for 3 min, followed by 40 cycles of 95°C denaturation for 10 s, and 60°C annealing/extension for 30 s. Relative *BMAL1* and *PER2* expression were determined by comparing *C*_*t*_ values of *BMAL1* and *PER2* to GAPDH (control) via 2^∧^ΔΔC_t_ method.^14^ Three biological replicates and three technical replicates per biological replicate were analyzed for each condition.

The Metacycle R package^15^ (https://cran.r-project.org/package=MetaCycle) was used to determine whether recordings were rhythmic in the circadian range. Default ranges were used for period (20-28 h) and the Bonferroni method was used to combine p-values from the three tests (Lomb-Scargle, ARSER, and JTK_Cycle).

### Lentiviral Transductions

Plasmids were generated and stably transfected as previously described.^16^ Briefly, 2.5 × 10^6^ HEK293T cells were seeded in 60 mm culture dishes and transiently transfected with 3 µg psPAX packaging plasmid, 2 µg pMD2G envelope plasmid, and 3 µg *BMAL1*:*luc* or *PER2*:*luc* reporter constructs using Lipofectamine3000 (ThermoScientific) according to manufacturer’s instructions. Lentiviral particles were harvested from supernatant and passed through a 45 µm filter. 9 mL lentivirus-containing supernatant was combined with 9 mL DMEM culture media (with all growth supplements) containing 10 µg/mL polybrene (Sigma). MCF7 (passage 11) and MDA-MB-231 (passage 16) cells were seeded in T25 culture flasks at 2 × 10^5^ cells/mL and incubated under standard conditions until 70-80% confluence was reached. Culture media was then removed, and 6 mL lentivirus-containing media added to each flask. After two days of infection, media was replaced with selection media (DMEM with all growth supplements plus 4 µg/mL puromycin) to obtain stably transfected cells. All cells were incubated at 37°C under 5% CO_2_ atmosphere.

### Bioluminescence Recording

Cells were seeded in 35 mm culture dishes at 2 × 10^5^ cells/mL and incubated for 2-3 days. Then, cells were starved and serum shocked as above. After 2 h synchronization, media was replaced with recording media consisting of DMEM (Sigma), 1% HEPES, 1% penicillin/streptomycin, 1% sodium pyruvate, 5% FBS, and 0.5 mM luciferin (ThermoScientific). Dishes were sealed with 40 mm sterile cover glass using silicon vacuum grease and subjected to continuous monitoring using a LumiCycle 32 Instrument (Actimetrics) at 36.5°C. Cells were not used at higher than passage 16 for MCF7 cells and passage 20 for MDA-MB-231.

Bioluminescence recordings were pre-processed to remove the initial transient (24 hours) and spikes. Each spike (any value more than 1/3^rd^ of the data range higher than that of the value at the preceding time step) was replaced with the average of the values of the preceding and succeeding time steps. To determine if a recording was rhythmic, we applied an FFT-based test^17^ to the time-series after removing a quadratic trend. Rhythmic time-series were de-noised and de-trended using the maximum overlap discrete wavelet transform (12-tap symmlet) with reflective boundary condition (WMTSA Matlab package written by Charles Cornish implementing previously published methods^18^), retaining the signal in the circadian band (periods of 21.33-42.67 h). Periods of the smoothed data were estimated using a continuous wavelet analysis (WAVOS Matlab package^19^), choosing the Morlet wavelet, excluding edge data, with a tuning parameter eta of 5, looking for a period within the range of 6-60 hours.

### Western Blot

MCF7 and MDA-MB-231 cells were cultured until confluent, and harvested using RIPA buffer (150 mM NaCl, 1% Triton X-100, 0.5% sodium deoxycholate, 0.1% sodium dodecyl sulfate (SDS), 50 mM Tris, pH 8.0, and freshly prepared 1% Halt™ Protease and Phosphatase Inhibitor (ThermoFisher)). Cell lysates were agitated for 30 min at 4°C and centrifuged at rt at 12,000 rpm for 20 min. Total cellular protein concentration was quantified via BCA Protein Assay Kit (ThermoFisher). 10 µg protein per sample was separated via 8% SDS polyacrylamide gel and transferred to PVDF membrane (ThermoFisher). Membrane was blocked with 5% (w/v) bovine serum albumin (BSA) in TBST (150 mM NaCl, 20 mM Tris-HCl, 0.1% Tween 20), incubated with primary antibodies against BMAL1 (Cell Signaling, cat no. 14020S), PER2, and GAPDH (Proteintech, cat no. 20359-1-AP and 20359-1-AP, respectively) overnight at 4°C, washed with TBST, and incubated with horseradish peroxidase-conjugated goat-anti-rabbit IgG secondary antibody (ThermoFisher) for 1 h at rt. Immunoblots were imaged using an enhanced chemiluminescence system (ECL; ThermoFisher) and a G:Box iChemi XT imaging system (GeneSys). Band intensities were analyzed via ImageJ.

## Results and Discussion

To confirm prior studies,^6,7,10^ we obtained RT-qPCR data tracking *BMAL1* and *PER2* in MCF7 and MDA-MB-231 cells. While both transcripts were present and detectable (**Fig. 1**), oscillations were not rhythmic in the circadian range (p>0.05 using Metacycle). To evaluate circadian rhythms in a more detailed manner to reveal subtle changes, we elected to employ luciferase reporters. This method has been used to provide high resolution oscillations in a variety of models,^1,11^ but not in breast cancer.

**Figure 1.**
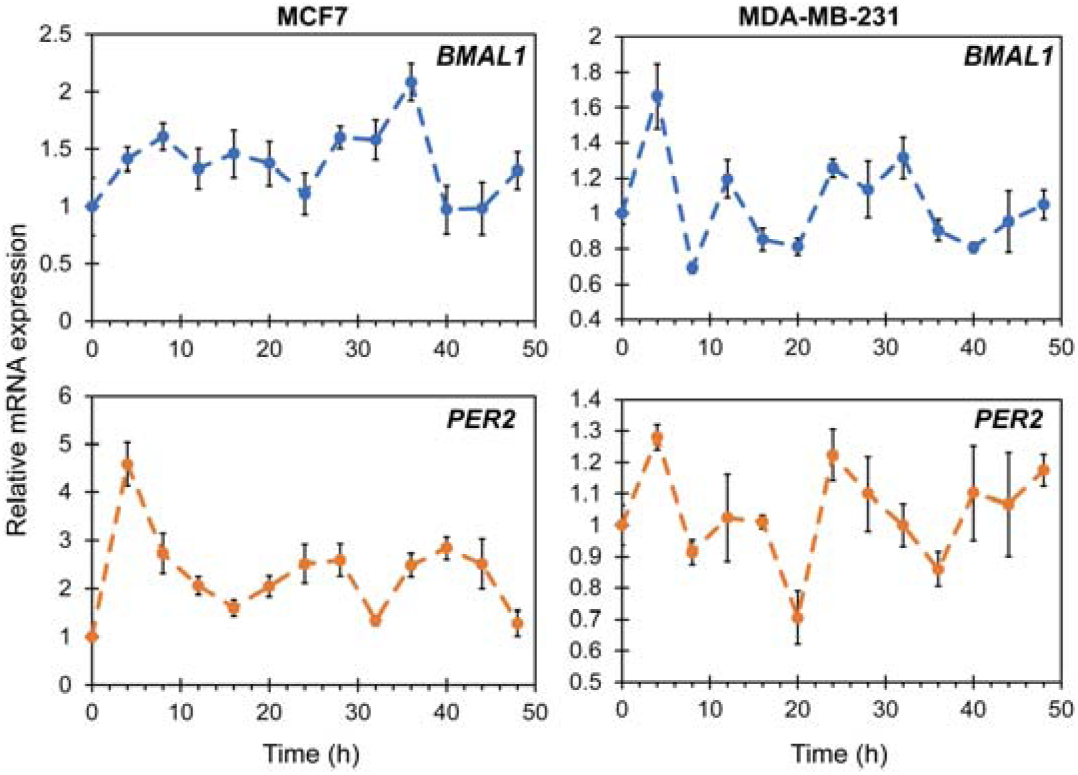
Quantitative RT-qPCR of *BMAL1* and *PER2* expression in MCF7 (left) and MDA-MB-231 (right) cells. Oscillations do not occur in a rhythmic manner (Metacycle rhythmicity test p > 0.05). Each point represents the average of three biological replicates, with three technical replicates each. Error bars represent relative errors.

To test our hypothesis that RT-qPCR assessments lack sufficient sensitivity to determine the presence or absence of circadian rhythms, we separately stably transfected *BMAL1:luc* and *PER2:luc* into MCF7 cells (**Fig. S1A** and **B**). Subsequent luminometry experiments revealed that circadian oscillations of both *BMAL1:luc* and *PER2:luc* persisted with a typical antiphase relationship (**Fig. 2** and **S2**). The trend in the raw bioluminescence time-series was negatively sloped and curving, and therefore appeared to be quadratic or exponential. Removal of a quadratic (**Fig. 2A** and **2B**) or exponential (**Fig. S3**) trend leads to qualitatively similar curves. An FFT-based test confirmed rhythmicity (p<0.05) for all experiments.^17^

**Figure 2.**
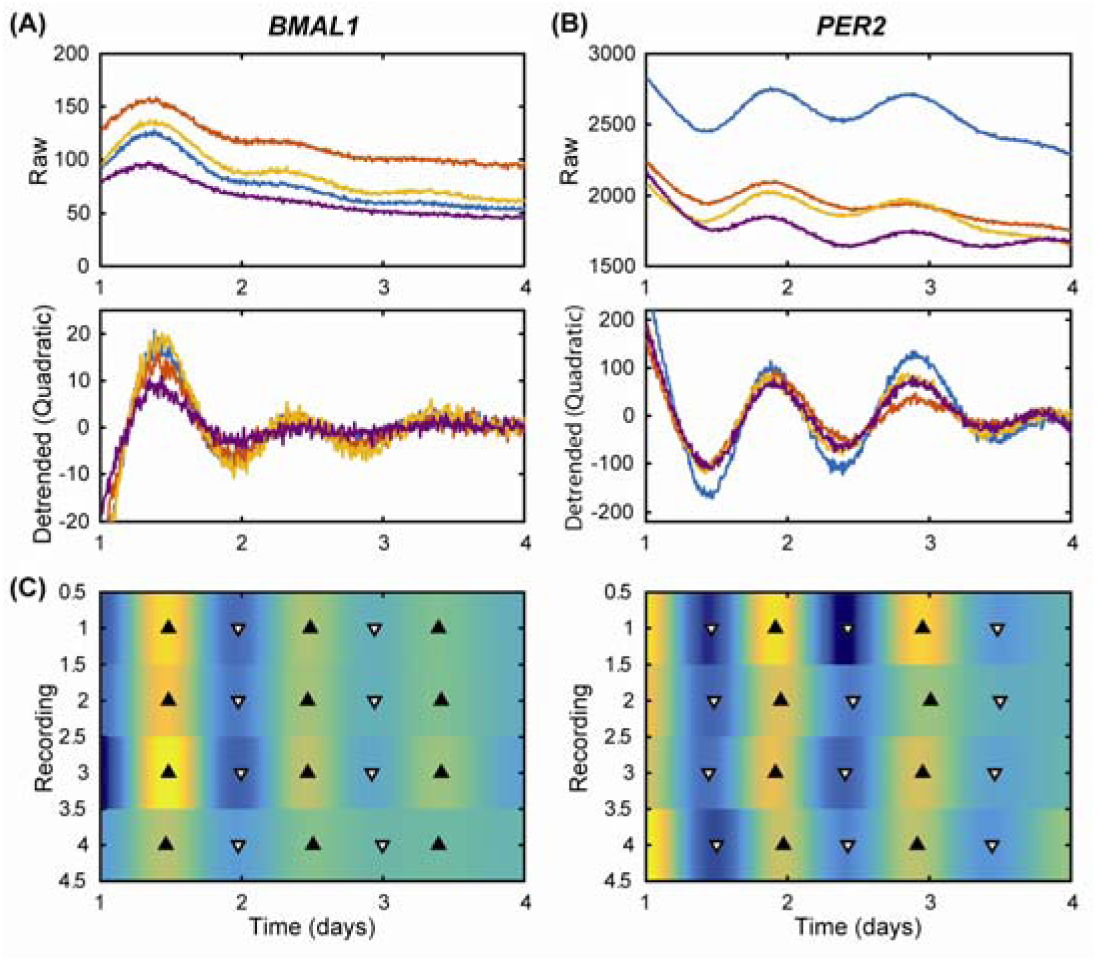
MCF7-*BMAL1:luc* **(A)** and *PER2:luc* **(B)** recordings have damped, anti-phase rhythms. Shown are time-series in raw form (top), with a quadratic trend removed (middle), and with the trend and noise removed by discrete wavelet analysis shown as a heat map **(C)**. The heat map indicates luminescence for each recording (N =4) with peaks (black triangles) and troughs (white triangles). Rhythms are clear, but lose amplitude over time. As expected in a functioning clock, peak times for *PER2:luc* are consistent and are 12 h out of phase with those of *BMAL1:luc*. Results from additional experiments are in **Fig. S2**.

To further quantify rhythms, we used continuous wavelet analysis to estimate the period of each recording over time, and computed the mean over time to yield a single period estimate for each recording. For data shown in **Fig. 2**, period estimate ranges were 22.5-23.2 h for *BMAL1:luc* and 22.1-23.6 h for *PER2:luc.* For the eight additional recordings (**Fig. S2**) the ranges are wider (22.4-25.2 h for *BMAL1:luc* and 18.7-23.3 h for *PER2:luc*). All period data are reported in **Table S1** and **S2**. These results stand in contrast with RT-qPCR analyses, which suggest that circadian clock gene expression is arrhythmic in breast cancer cells.

To test our hypothesis that with increasing breast tumor grade/aggressiveness, cellular circadian rhythms are increasingly disrupted, we stably transfected *BMAL1:luc* and *PER2:luc* separately into triple negative/basal MDA-MB-231 cells (**Fig. S1C** and **S1D**). The luminometry data from these highly aggressive cells revealed no visually detectable oscillations (**Fig. 3, Fig. S4**), even after removal of either a quadratic (**Fig. 3**) or exponential (**Fig. S5**) trend. Very few statistically significant oscillations were observed (FFT-based test for rhythmicity yielded p<0.05 for only 1 of 12 *BMAL1:luc* and 3 of 12 *PER2:luc* recordings).^17^ In addition to assessing transcriptional activities of *BMAL1* and *PER2* in both cancer cell lines, we examined overall protein expression levels (**Fig. 4**). Our western blot data corroborates the presence of BMAL1 and PER2, and indicates diminished levels of both with increased tumor grade/aggressiveness from luminal A to basal.

The subtype-defining molecular differences between MCF7 and MDA-MB-231 cell lines are presence of estrogen receptor (ER) and progesterone receptor (PR) in MCF7 cells. Studies have shown that cellular treatment with 17β-estradiol (E2) and progesterone (P4), the ligands for ER and PR, can affect circadian oscillations in normal cells expressing these receptors.^20^ It has also been shown that E2 treatment can enhance expression of *BMAL1, CLOCK, PER1*, and *PER2*.^9,21,22^ We hypothesize that the presence of the ER-E2 pathway may contribute to the circadian oscillations present in MCF7 cells. Future studies will examine other ER-positive breast cancer cell lines (i.e. ZR-75-1) to evaluate whether results obtained here remain consistent.

**Figure 3.**
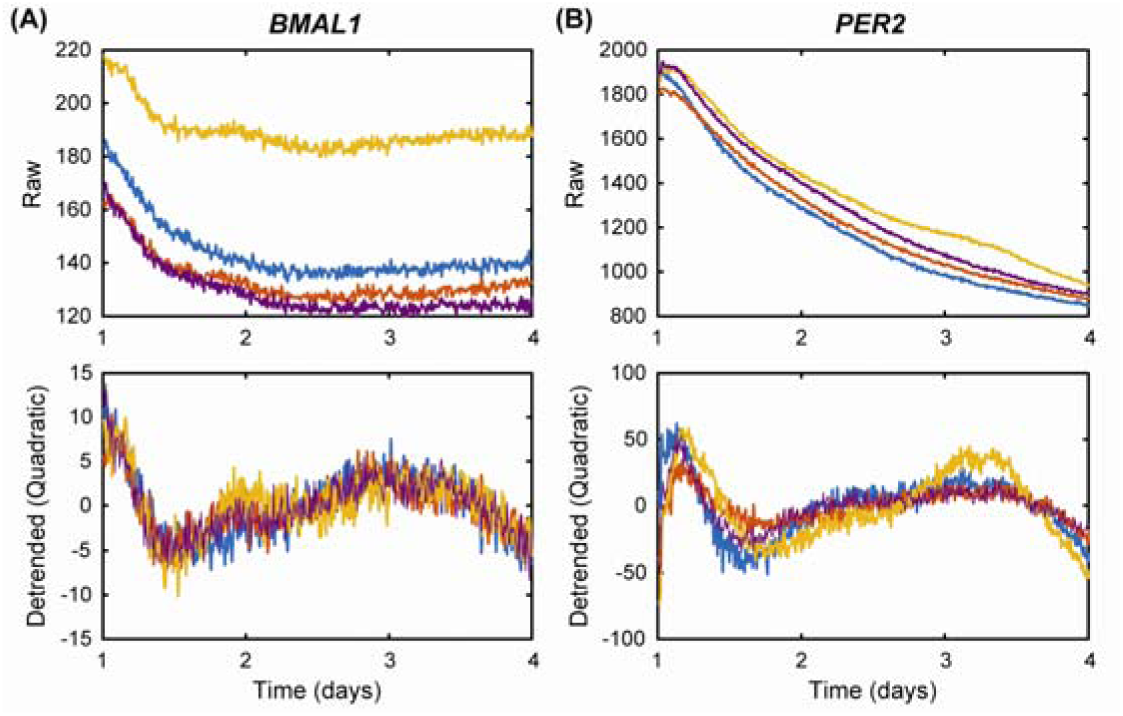
MDA-MB-231-*BMAL1:luc* **(A)** and *PER2:luc* **(B)** recordings reveal no circadian rhythms. Shown are time-series in raw form (top) and with a quadratic trend removed (bottom). Visual inspection and an FFT-based test for rhythmicity confirm there are no circadian rhythms in these recordings. Results from additional experiments are in **Fig. S3.**

**Figure 4.**
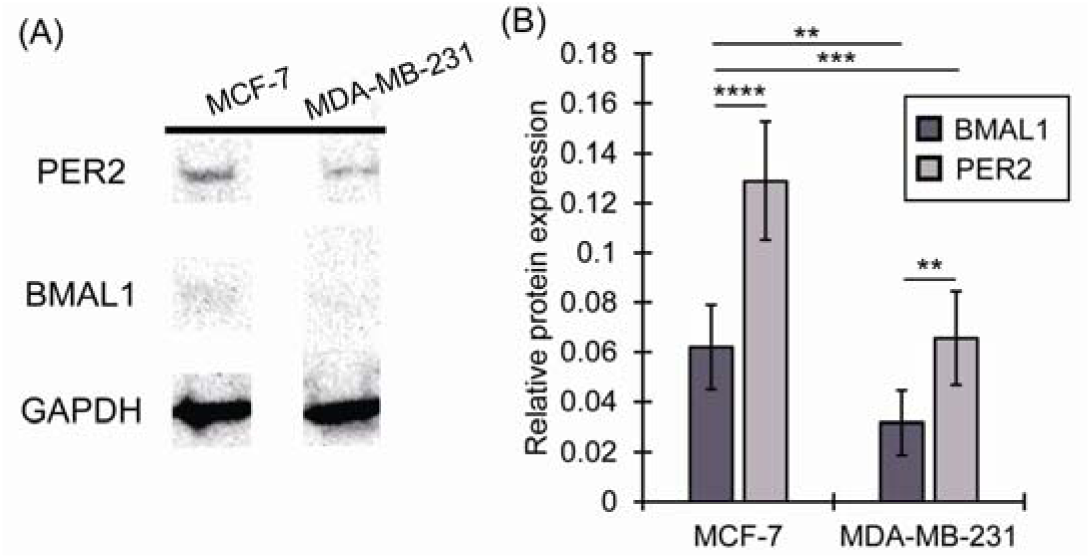
Expression of core clock proteins BMAL1 and PER2 in MCF7 and MDA-MB-231 cells. Representative western blot showing PER2 and BMAL1 expression levels in MCF7 and MDA-MB-231 cells **(A)**. Western blots (two biological replicates with three technical replicates each) were assessed via densitometry **(B)**. Protein expression of BMAL1 and PER2 was normalized to GAPDH; paired student T-tests were used to calculate significance (**p<0.01, ***p<0.001,****p<0.0001).

Another difference between MCF7 and MDA-MB-231 cells occurs in terms of epidermal growth factor receptor (EGFR) status.^23^ MDA-MB-231 cells express EGFR to a greater degree than do MCF7 cells. Following activation via EGF, the EGFR cascade’s downstream pathways, including Ras/MAPK and PI3K/mTOR,^24^ have been shown to disrupt circadian rhythms via period alteration or clock gene suppression.^25,26^ This may be a cause of the arrhythmic oscillations observed in MDA-MB-231 cells. In the future, the role of EGFR in the circadian network should be further assessed.

Based on the data obtained, we conclude that RT-qPCR assessments are not sufficient to track low amplitude, subtle circadian oscillations in cancer cells. Additionally, we show that the clock is not arrhythmic (or “broken”) in all breast cancers, as had previously been posited. Furthermore, via assessment of MDA-MB-231 cells, we provide initial evidence that circadian rhythms of both positive (*BMAL1*) and negative (*PER2*) components of the clock are disrupted with increasing tumor grade. Our results indicate that studies in additional representatives of breast cancer subtypes is highly merited.

## Acknowledgements

We acknowledge Prof. Tanya Leise (Mathematics and Statistics, Amherst College) and Prof. D. Joseph Jerry (Veterinary and Animal Sciences, UMass Amherst) for helpful comments on the manuscript. We thank Prof. Jungwoo Lee (Chemical Engineering, UMass Amherst) for plate reader and Prof. Elizabeth Vierling (Biochemistry and Molecular Biology, UMass Amherst) for gel documentation system access.

