## Supporting Information for "Circadian Oscillations Persist in Low Malignancy Breast Cancer Cells"

**RNA Extraction and cDNA Synthesis**

Following synchronization of cells by serum shock, cells were washed with PBS and returned to starvation conditions. Cells were harvested with the first time point (T=0) taken prior to serum shock, and every 4 h thereafter for 48 h. Total RNA was extracted via TRIzol Reagent (Gibco) according to the manufacturer’s instructions. 1 mL TRIzol Reagent was added to lyse the cells. Cell lysates were incubated at rt for 5 min to allow complete dissociation of nucleoprotein complexes. After addition of 200 μL chloroform per 1 mL TRIzol, samples were shaken vigorously by hand for 15 s and incubated at rt for 3 min. Samples were then centrifuged at 12,000 x g for 15 min at 4°C and the upper phase containing RNA was separated. The RNA samples were further purified via PureLink RNA kit (Ambion) according to the manufacturer’s instructions. Total RNA concentration was determined via Nanodrop UV/Vis (Thermo Fisher Scientific). 1 μg of total RNA was reverse-transcribed to cDNA using 50 μM random hexamers, 40 U/μL RNaseOut, 10 mM dNTPs, and 200 U/μL SuperScript IV Reverse Transcriptase (Thermo Fisher Scientific).

**Quantitative real-time PCR (RT-qPCR)**

The following primer sequences were used: *GAPDH* forward (5'- CTT CTT TTG CGT CGC CAG CC-3'), reverse (5'-ATT CCG TTG ACT CCG ACC TTC-3'); *BMAL1* forward (5’- CTA CGC TAG AGG GCT TCC TG-3’), reverse (5’- CTT TTC AGG CGG TCA GCT TC-3’); *PER2* forward (5’- TGT CCC AGG TGG AGA GTG GT-3’), reverse (5’- TGT CAC CGC AGT TCA AAC GAG-3’).

**Luciferase assays**

Cells were seeded in 24-well plates at a density of 2x10^5^ cells/mL and incubated for approximately 1-3 days. After cells reached 100% confluence, they were lysed using the Luciferase assay system (Promega) according to the manufacturer’s instructions. Luminescence was measured using a Synergy H1 microplate reader.

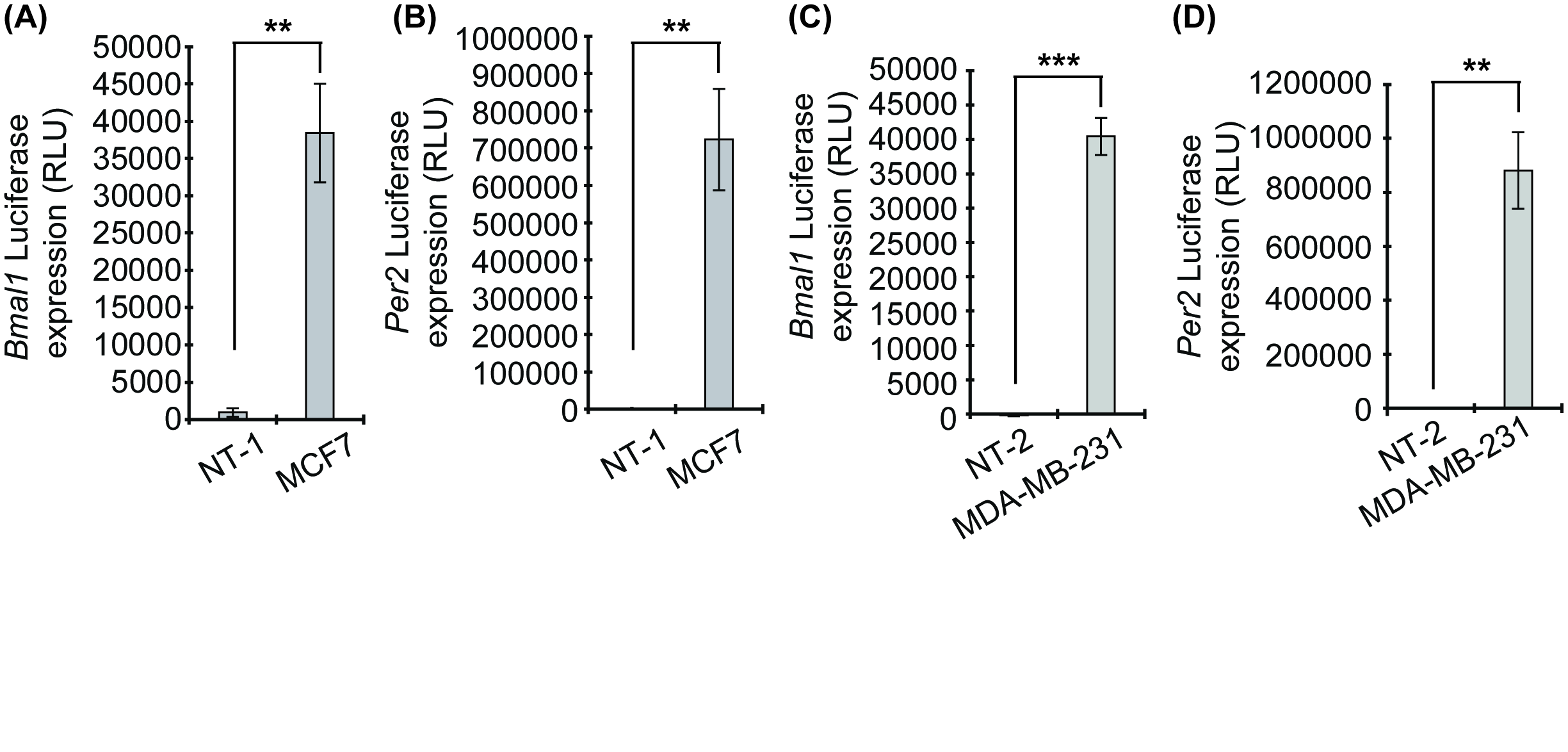

**Figure S1.** Luciferase assay data obtained for the stable transfections of BMAL1:luc **(A, C)** and PER2:luc **(B, D)** in MCF7 and MDA-MB-231 cells. Each condition has three technical replicates (n = 3). Paired student T-tests were used to calculate significance between conditions (**p<0.001,***p<0.0001). NT-1 = non-transfected MCF7 cells; NT-2 = non-transfected MDA-MB-231 cells.

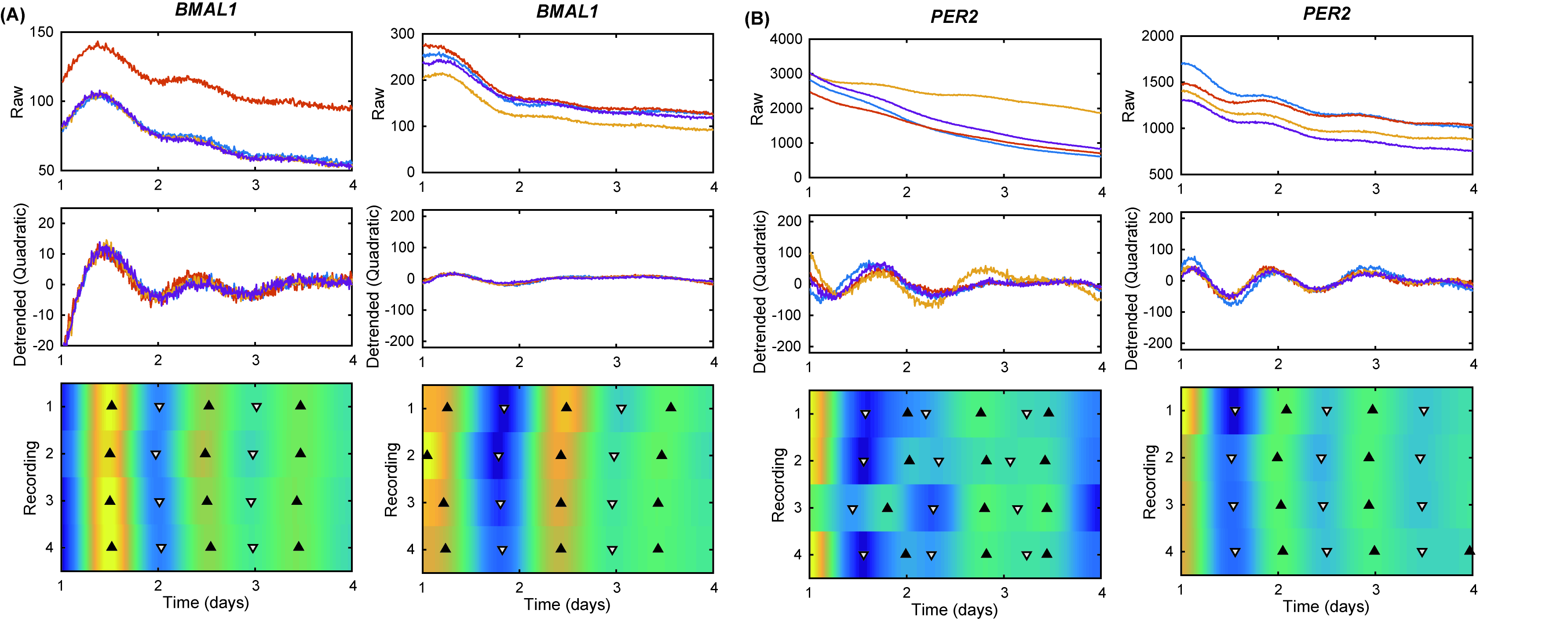

**Figure S2.** *BMAL1:luc* **(A)** and *PER2:luc* **(B)** recordings in MCF7 cells have damped, anti-phase rhythms. Shown are results from four additional experiments (2 each per reporter), including the time-series in raw form (top), with a quadratic trend removed (middle), and with the trend and noise removed by a discrete wavelet analysis as a heat map (bottom). The heat map indicates the luminescence for each recording (N = 4) with peaks (black triangles) and troughs (white triangles). Rhythms are clear, but lose amplitude. As expected in a functioning clock, the peak times for *PER2:luc* are consistent and 12 h out of phase with those of *BMAL1:luc*.

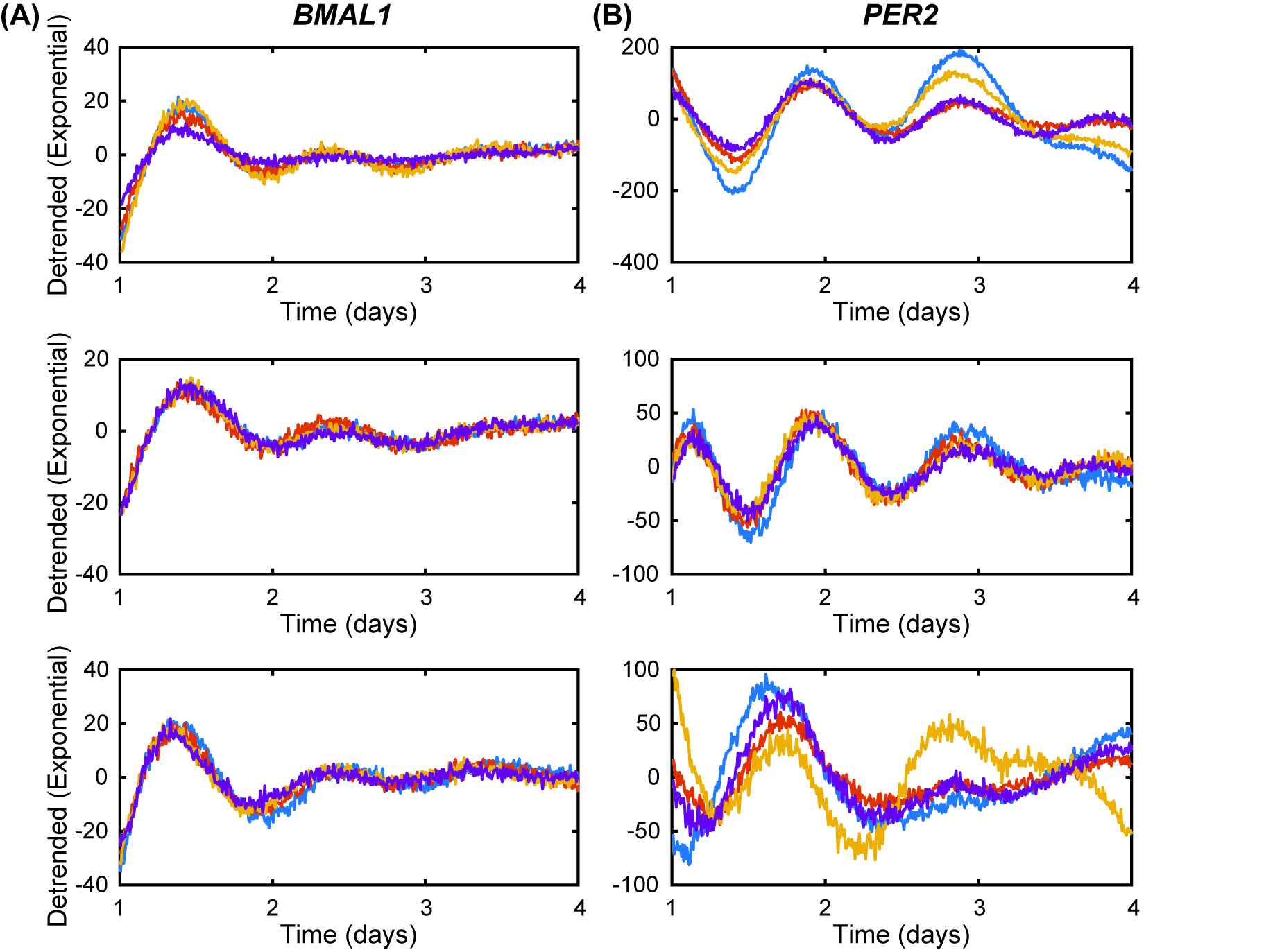

**Figure S3.** Bioluminescence recordings of MCF7 with *BMAL1:luc* **(A)** and *PER2:luc* **(B)** reporters after removing exponential trends. Shown are the recordings for 3 separate experiments (N = 4 in each).

|  | Replicate 1 | Replicate 2 | Replicate 3 | Replicate 4 |
| --- | --- | --- | --- | --- |
| Experiment 1 | 22.5 h | 22.5 h | 22.2 h | 23.2 h |
| Experiment 2 | 22.7 h | 22.8 h | 22.4 h | 22.5 h |
| Experiment 3 | 25.2 h | 25.2 h | 25.2 h | 25.2 h |

**Table S1.** Period estimates from Bioluminescence recordings of MCF7 with *BMAL1:luc* reporter. Shown are the estimates for each of 3 experiments (N=4 in each). Periods were estimated using a continuous wavelet transform on recordings that were de-noised and de-trended by a discrete wavelet transform. The continuous wavelet period was estimated for each time step and we report the average (taken over time).

|  | Replicate 1 | Replicate 2 | Replicate 3 | Replicate 4 |
| --- | --- | --- | --- | --- |
| Experiment 1 | 23.5 h | 23.6 h | 23.5 h | 22.1 h |
| Experiment 2 | 22.6 h | 22.2 h | 22.2 h | 23.2 h |
| Experiment 3 | 19.9 h | 23.3 h | 18.8 h | 18.7 h |

**Table S2.** Period estimates from Bioluminescence recordings of MCF7 with *PER2:luc* reporter. Shown are the estimates for each of 3 experiments (N=4 in each). Periods were estimated using a continuous wavelet transform on recordings that were de-noised and de-trended by a discrete wavelet transform. The continuous wavelet period was estimated for each time step and we report the average (taken over time).

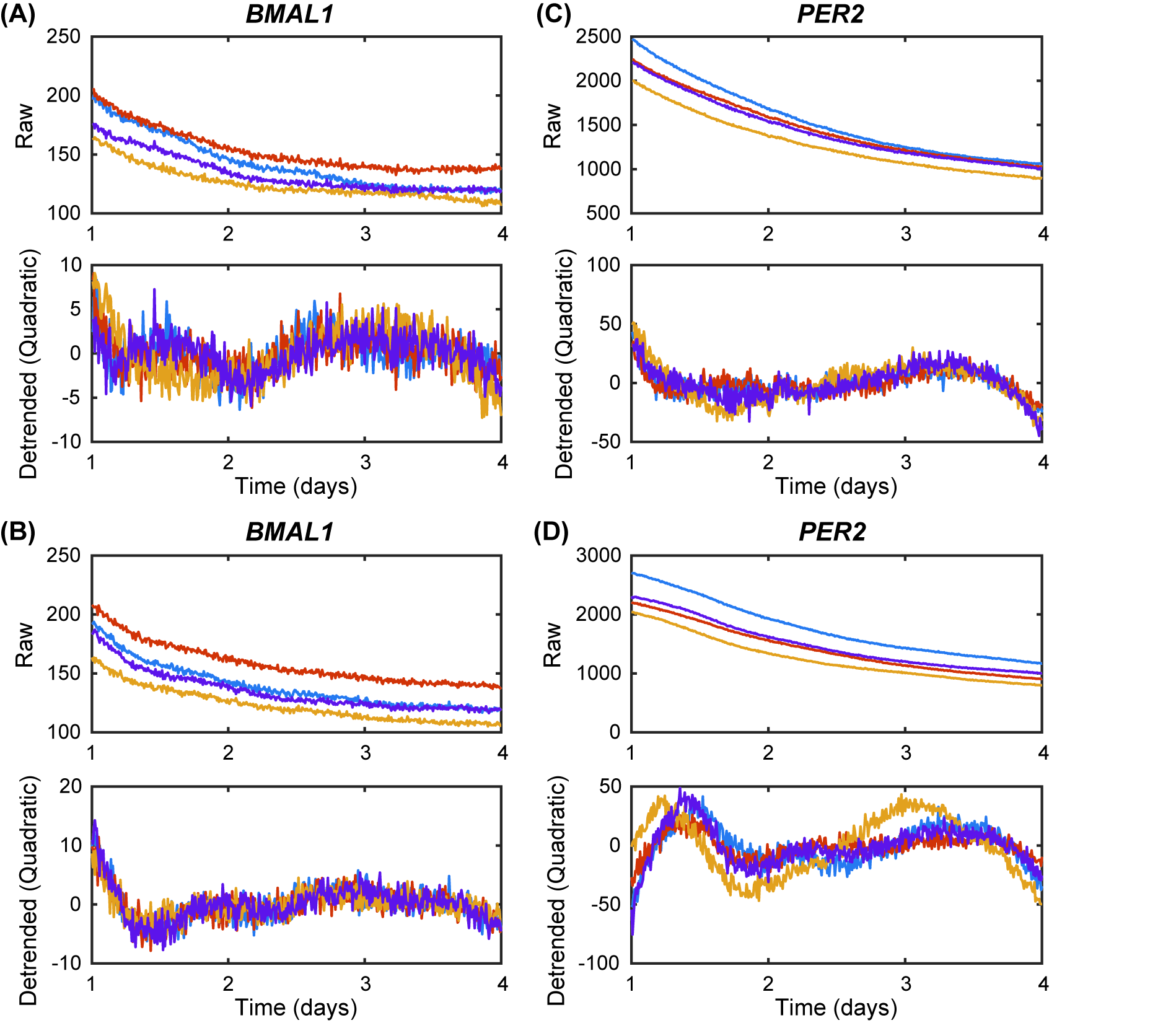

**Figure S4.** MDA-MB-231-*BMAL1:luc* **(A, B)** and *PER2:luc* **(C, D)** recordings (N = 4 in each) have no clear circadian rhythms. Shown are the time-series in raw form (rows 1 and 3) and with a quadratic trend removed (rows 2 and 4). An FFT-based test for rhythmicity found only 1 *BMAL1:luc* recording and only 3 *PER2:luc* recordings rhythmic.

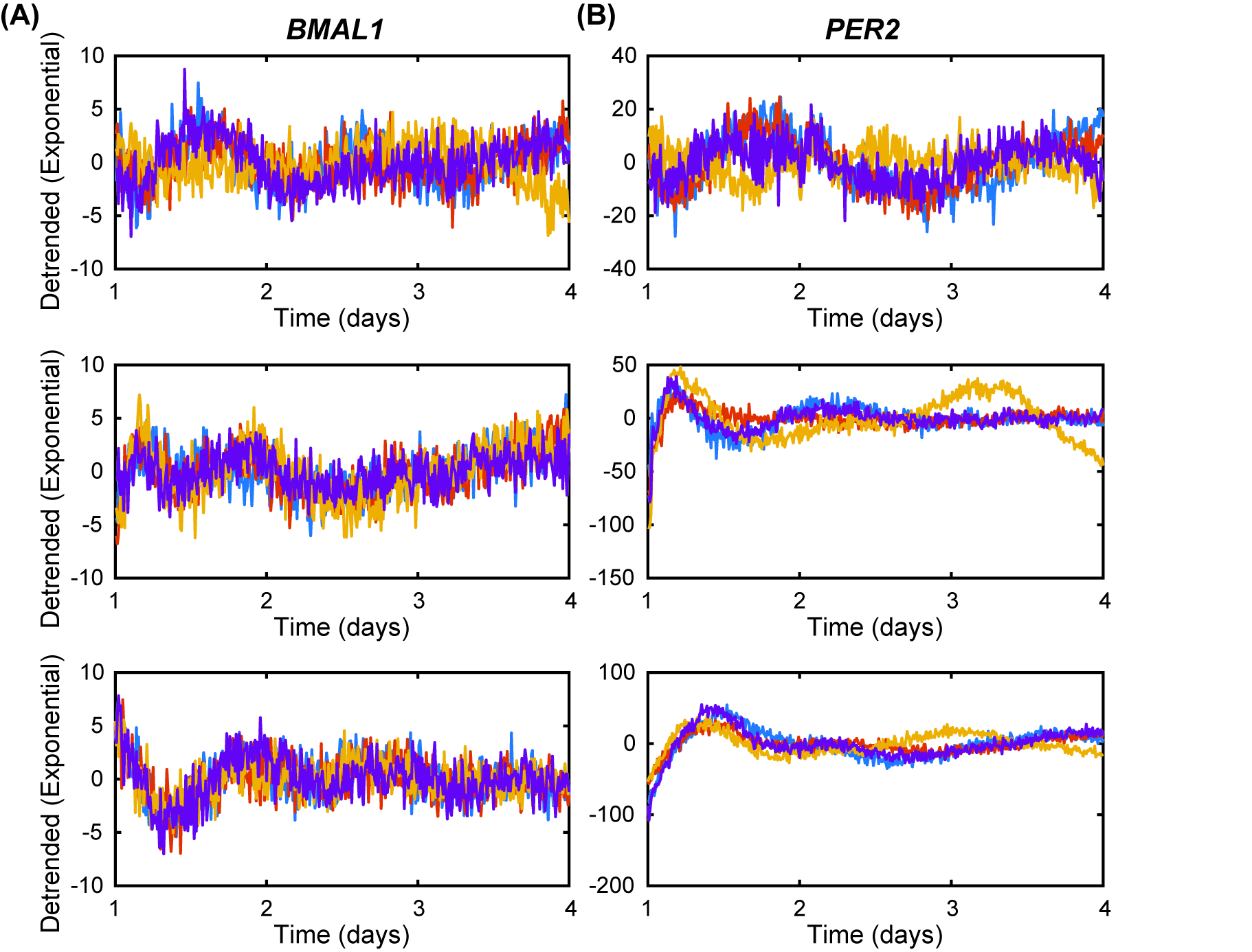

**Figure S5.** Bioluminescence recordings of MDA-MB-231 with *BMAL1:luc* **(A)** *and PER2:luc* ***(B)*** reporters. Shown are the recordings for each of 3 experiments (N=4 in each) after removing exponential trends.
